# Collaborative environmental DNA sampling from petal surfaces of flowering cherry *Cerasus ×- yedoensis* 'Somei-yoshino' across the Japanese archipelago

**DOI:** 10.1101/165522

**Authors:** Tazro Ohta, Takeshi Kawashima, Natsuko O. Shinozaki, Akito Dobashi, Satoshi Hiraoka, Tatsuhiko Hoshino, Keiichi Kanno, Takafumi Kataoka, Shuichi Kawashima, Motomu Matsui, Wataru Nemoto, Suguru Nishijima, Natsuki Suganuma, Haruo Suzuki, Y-h. Taguchi, Yoichi Takenaka, Yosuke Tanigawa, Momoka Tsuneyoshi, Kazutoshi Yoshitake, Yukuto Sato, Riu Yamashita, Kazuharu Arakawa, Wataru Iwasaki

## Abstract

Recent studies have shown that environmental DNA is found almost everywhere. Flower petal surfaces are an attractive tissue to use for investigation of the dispersal of environmental DNA in nature as they are isolated from the external environment until the bud opens and only then can the petal surface accumulate environmental DNA. Here, we performed a crowdsourced experiment, the “Ohanami Project”, to obtain environmental DNA samples from petal surfaces of *Cerasus × yedoensis* ‘Somei-yoshino’ across the Japanese archipelago during spring 2015. *C. × yedoensis* is the most popular garden cherry species in Japan and clones of this cultivar bloom simultaneously every spring. Data collection spanned almost every prefecture and totaled 577 DNA samples from 149 collaborators. Preliminary amplicon-sequencing analysis showed the rapid attachment of environmental DNA onto the petal surfaces. Notably, we found DNA of other common plant species in samples obtained across a wide distribution; this DNA likely originated from pollen of the Japanese cedar. Our analysis supports our belief that petal surfaces after blossoming are a promising target to reveal the dynamics of environmental DNA in nature. The success of our experiment also shows that crowdsourced environmental DNA analyses have considerable value in ecological studies.

## INTRODUCTION

Recent research has shown that environmental DNA sequences are present everywhere (Rosario and Breitbart 2011). Although environmental DNA can be sampled from a wide range of targets, the petal surface of flowers is an attractive source for investigation as it enables analysis of the dispersal of environmental DNA into the emerging niche of a newly-opened flower.

The ornamental cultivar ‘Somei-yoshino’ (*Cerasus × yedoensis*) is the most widely cultivated cherry tree in Japan (Lindstrom 2007, Shirahata 2000), with the exception of Okinawa prefecture (Iketani et al. 2007). In part, because *C. × yedoensis* is propagated through grafting (and therefore cloned from a single tree), it blooms synchronously within a single climate (Innan et al. 1995, Kato et al. 2012). The pale-rose five-petal flowers are considered a symbol of spring in Japan, and “Ohanami” (flower viewing) parties are held annually (Fig. 1A). Upon bud opening, *C. × yedoensis* petal surfaces exhibit a *petal effect*, an adhesive quality that causes the capture of environmental DNA in the form of microbes or pollen deposited by pollinators or the wind (Feng et al. 2008). Differences in the environmental DNA on petal surfaces of cloned plants may reflect differences in the interaction of the plants with their environments. To date, however, there is very little data on environmental DNA samples from cloned plants to investigate differences among geographical locations.

**Fig. 1.**
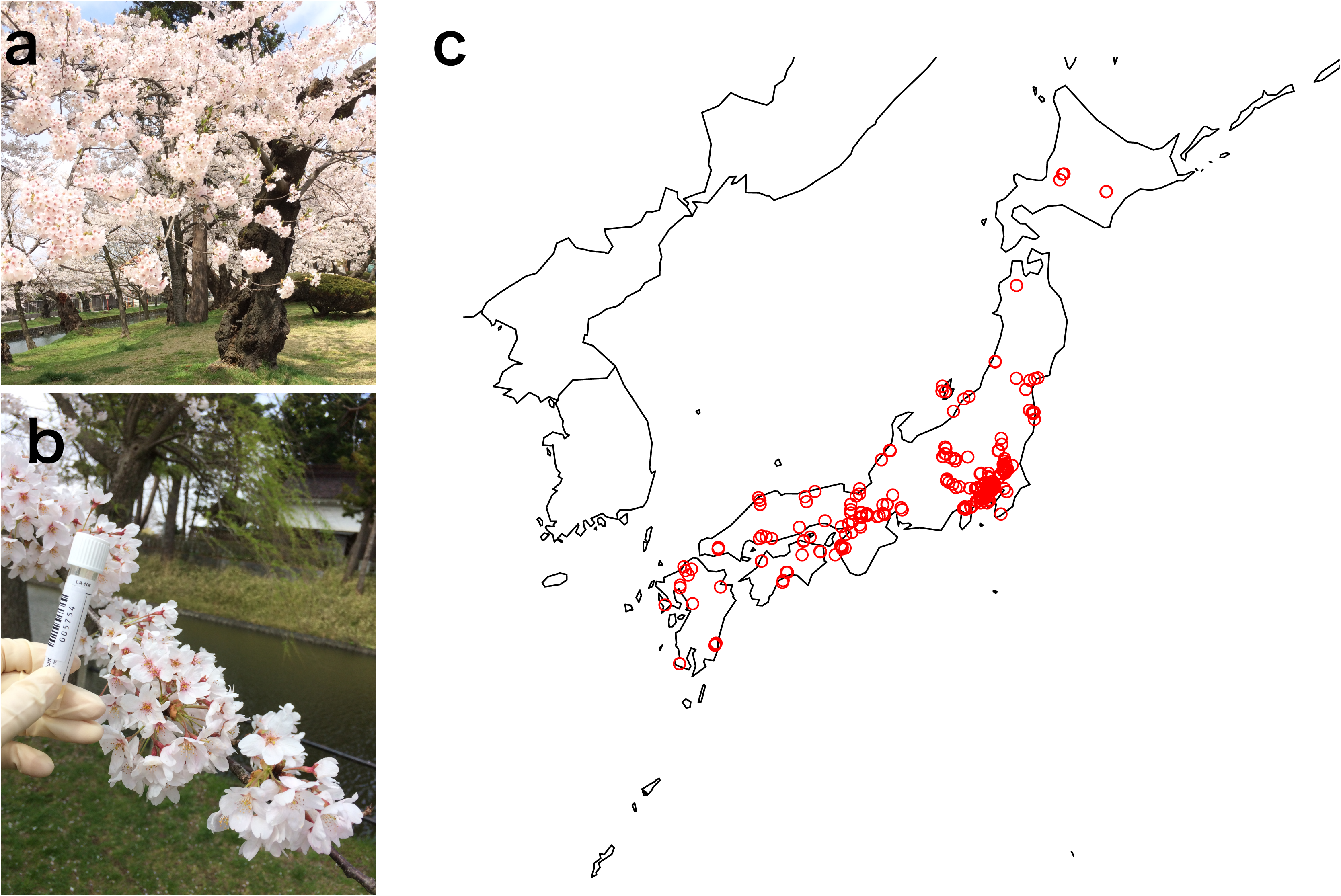
*Cerasus × yedoensis* and the “Ohanami Project.” **a** Photographs of a typical landscape at a sampling site during the cherry-blossom season in Japan and **b** environmental DNA sampling of *C. × yedoensis* petal surfaces (sample id 005789 at Tsuruoka, Yamagata prefecture). The corymbose inflorescence has three to five flowers arranged on a stem, and each corolla has five petals attached to the hypanthium rim. **c** A geographical map of the 123 sampling sites. Latitudinal and longitudinal coordinates of the sites are shown in the Supplementary File 2.

Our goals in this study were two-fold. First, we aimed to elucidate the origins of environmental DNA present on flower petals. Second, we aimed to examine the effectiveness of a crowdsourcing project (Howe 2006) for rapid and large-scale sampling. In investigations into environmental DNA, it is critical to collect sufficiently large sample from multiple locations, while retaining sample quality. A previous study used a crowdsourcing approach to solve the problem of sampling from many geographically separate locations (MetaSUB International Consortium, 2016). This method potentially offers an effective approach for sample collection across a large area and over a very short interval.

Our “Ohanami Project” took place during spring 2015 and targeted environmental DNA on petals of newly-opened flowers of *C. × yedoensis*. In this report, we present the results of a crowdsourcing experiment to sample DNA from the petal surfaces of *C. × yedoensis* and the subsequent amplicon sequencing analysis to identify the sources of the foreign DNAs.

## Materials and methods

### Crowdsourced environmental DNA sampling of C. × yedoensis petal surfaces

Sampling was conducted from March 17 to May 5, 2015. A sampling kit containing a disposable mask, disposable gloves, and barcode label was sent to 149 collaborators across Japan. We chose Puritan(r) Opti-Swab(r) Liquid Amies Collection & Transport System LA-106 as the swab kit. This kit contains a buffer composed of 3.0 g sodium chloride, 0.2 g monopotassium phosphate, 1.2 g disodium phosphate, 0.2 g potassium chloride, 1.0 g sodium thioglycolate, 0.1 g calcium chloride, and 0.1 g magnesium chloride per liter. To ensure the sampling quality of the crowdsourced samples, we designed and attached a detailed sampling protocol (Supplementary File 1). Each collaborator followed this protocol to sample environmental DNA from *C. × yedoensis* petal surfaces in their neighborhood, swabbing more than 10 flowers per tree; masks and gloves were worn to avoid contamination by the collector (Fig. 1B). Immediately upon sampling, collaborators were instructed to affix the barcode label on the collection tube and take a photograph with their mobile phone; these steps were intended to minimize handling errors and to record locations through the Global Positioning System. We obtained 577 samples from 123 sites in 103 cities, and nearly every prefecture was covered, from Sapporo City, Hokkaido Prefecture (northernmost) to Kagoshima City, Kagoshima Prefecture (southernmost) (Fig. 1C and Online Resource 2). Just under half the samples (281, 48.7 %) were collected from March 30 to April 4, corresponding to the official 2015 cherry-blossom season reported by the Japan Meteorological Agency (2008). Twenty samples that were collected from other cherry species were excluded from the subsequent analyses: four were from *Cerasus jamasakura* in Abashiri, Hokkaido and 16 from other *Cerasus* sp. in Mishima, Shizuoka. Samples were stored at -20°C until refrigerated delivery to the sequencing center.

### DNA extraction and amplicon library preparation

. Each sample was centrifuged at 20,000 *g* for 10 min, and 150 μL of the resultant supernatant was boiled at 99°C for 15 min before immediately chilling on ice (Yamagishi et al. 2016). The sample was then centrifuged again (10,000 g for 15 min) to extract template DNA from the supernatant. PCR amplification was performed targeting the V4 hypervariable region of bacterial, mitochondrial, and chloroplast 16S ribosomal RNA (Kozich 2013). The initial PCR reaction volume was 10 μL, containing 1 μL template DNA, 1 μL 10× PCR buffer, 0.8 μL 10 mM dNTP mix, 0.25 μL each of 10 μM forward and reverse primers, 0.05 μL Ex Taq HS DNA Polymerase (Takara Bio, Kusatsu, Japan), and 6.65 μL of RNase-free water. Primers and index tag sequences are shown in Supplementary File 3-5. Thermocycling conditions comprised an initial denaturation at 94°C for 3 min; 40 cycles of denaturation at 94°C for 45 s, annealing at 50°C for 1 min, and extension at 72°C for 1.5 min; and a final extension at 72°C for 10 min.

A second PCR amplification was performed using a 20 μL reaction mixture: 1 μL of the first PCR product, 2 μL 10× PCR buffer, 1.6 μL 10 mM dNTP mix, 5 μL each of 1 μM PCR-F/R index primers, 0.1 μL Ex Taq HS DNA Polymerase, and 5.3 μL RNase-free water. The PCR thermocycling conditions were as follows: initial denaturation at 94°C for 3 min; 20 cycles of denaturation at 94°C for 45 s, annealing at 50°C for 1 min, and extension at 72°C for 1.5 min; and final extension at 72°C for 10 min. The length of the PCR product was 258 or 259 bp. After confirming successful amplification using a fragment analyzer (Advanced Analytical, Ankeny, USA), PCR products were size-selected using BluePippin (Sage Science, Beverly, USA). The products were then treated with ExoSAP-IT (Affymetrix, Santa Clara, USA) for enzymatic cleanup and purified using AMPure XP (Beckman Coulter, Brea, USA). A Qubit Fluorometer (ThermoFisher, Waltham, USA) was used to measure DNA concentration in pooled samples and the samples were adjusted to 10 pM.

### Amplicon sequencing

. Two runs of MiSeq sequencing (Illumina, San Diego, USA) were performed using the MiSeq Reagent kit v3 Box-2, following a previously reported protocol (Yamagishi et al. 2016). Read quality was checked by FastQC v0.10.1 (Andrews 2010). Using default settings, adapter sequences were removed using TagCleaner v0.16 (Schmieder et al. 2010), and paired-end reads were assembled using FLASH v1.2.7 (Magoč and Salzberg 2011). Reads were trimmed according to base call accuracy in DynamicTrim.pl v2.2 (Cox et al. 2010) and quality-filtered (i.e., reads containing Ns and with lengths ≤120 or ≥320 bp were removed).

### Taxonomic assignment

. Reads were taxonomically assigned using the Illumina BaseSpace System Metagenomics Workflow (Version 1.0.0.79, Culture = neutral option). Further classifications of reads grouped into Cyanobacteria were subsequently performed using BLASTN searches (Morgulis et al. 2008) against the chloroplast reference sequence dataset from Organelle Genome Resources at NCBI (Wolfsberg et al. 2001). Likewise, all reads grouped into Proteobacteria were classified with BLASTN searches against the mitochondrial reference sequence dataset from Organelle Genome Resources.

### Data availability

. Raw sequence data were deposited in the DDBJ public database under an accession number DRA005186.

## Results and discussion

In total, 28,594,048 sequence reads of 251 bp in length were obtained by MiSeq sequencing. After quality control, 5,606,853 reads remained for subsequent analyses. Initial taxonomic assignment of all reads identified various prokaryotic phyla (Fig. 2) in which Cyanobacteria and Proteobacteria dominated with 81% and 12%, respectively. According to BLASTN results, most of the reads assigned to Cyanobacteria and Proteobacteria were derived from chloroplasts and mitochondria of the host tree and other species. Specifically, the majority of reads exactly matched to chloroplast DNA from wild strawberry *Fragaria virginiana* (family Rosaceae) (Fig. Since both wild strawberry and flowering cherry belong to the Rosaceae family, it was expected that V4 hypervariable region sequences of the two species are identical. Thus, we compared the sequences of *F. virginiana* and the chloroplast genome sequence of *Prunus yedoensis*, retrieved from the NCBI GenBank (GenBank ID: KP760070), which was not included in the organelle database (Fig. 4). The alignment shows that there is no mismatch among the two reference sequences and the most abundant reads in our samples, which indicated that the reads assigned to *F. virginiana* by BLASTN were derived from the host plant. The second, third, and fourth most frequent reads were one-nucleotide-mismatch hits to *P. yedoensis* chloroplast DNA, with a sharp decrease in hits containing more than one mismatch (Fig. 3).

**Fig. 2.**
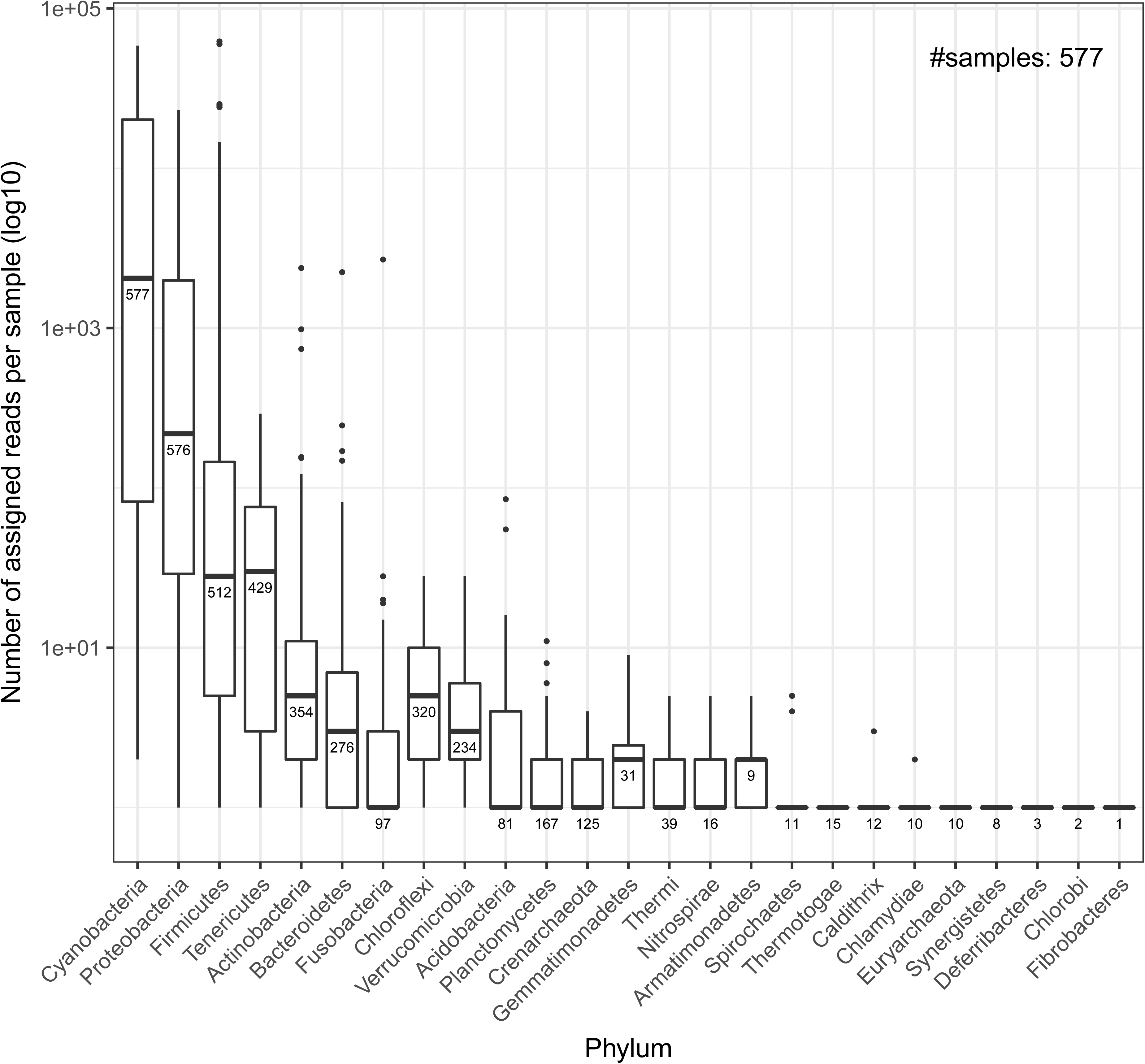
Phylum-level taxonomic abundance ratios estimated using the initial taxonomic assignment of 577 DNA samples from *C. × yedoensis* petals. The horizontal axis shows assigned phyla, and the vertical axis shows numbers of assigned reads per sample in the log10 scale. The bottom, central line, and top of the box plots represent the first, second, and third quartiles, respectively. Whiskers represent the first quartile -1.5 × Interquartile Range (IQR) and the third quartile +1.5 × IQR. The number below the central line of the box indicates number of samples that contained reads assigned to each phylum. The plot shows that the reads assigned to Cyanobacteria and Proteobacteria are found in almost all of the collected samples (577 and 576), and their numbers of reads per sample was larger than those of the other phyla.

**Fig. 3.**
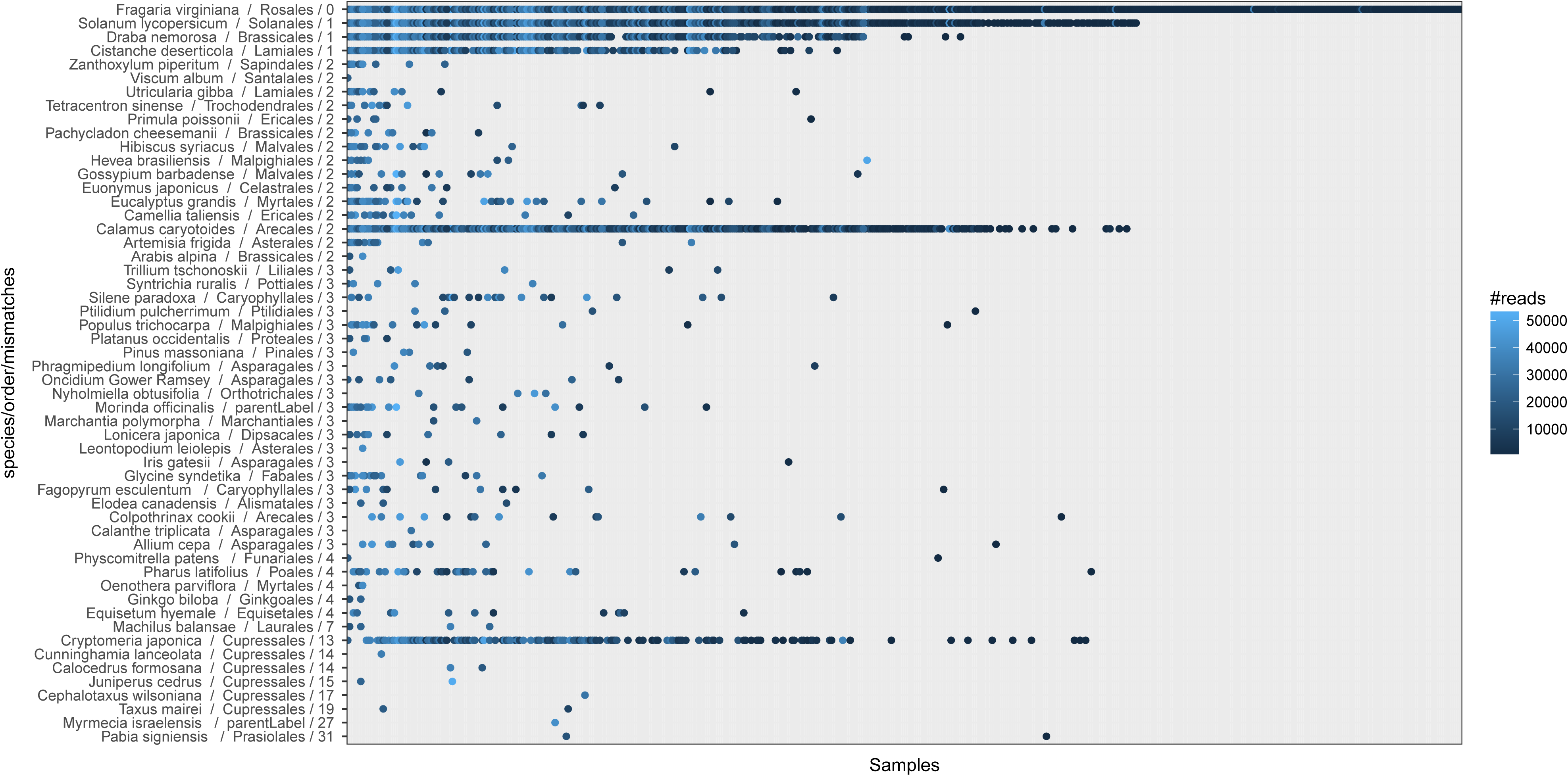
Results of sequence similarity searches against the Organelle Genome Resources for chloroplast genomes at NCBI. The vertical axis lists species names ordered by numbers of nucleotide mismatches against the sequence of *F. virginiana*, which sequence was assumed to be identical to that of the host plant. The horizontal axis represents 577 samples, sorted by numbers of species with hits against chloroplast DNA. The plot suggests that some species such as *Calamus caryotoides* and *Cryptomeria japonica* were frequently found in different samples, or in various locations.

**Fig. 4.**
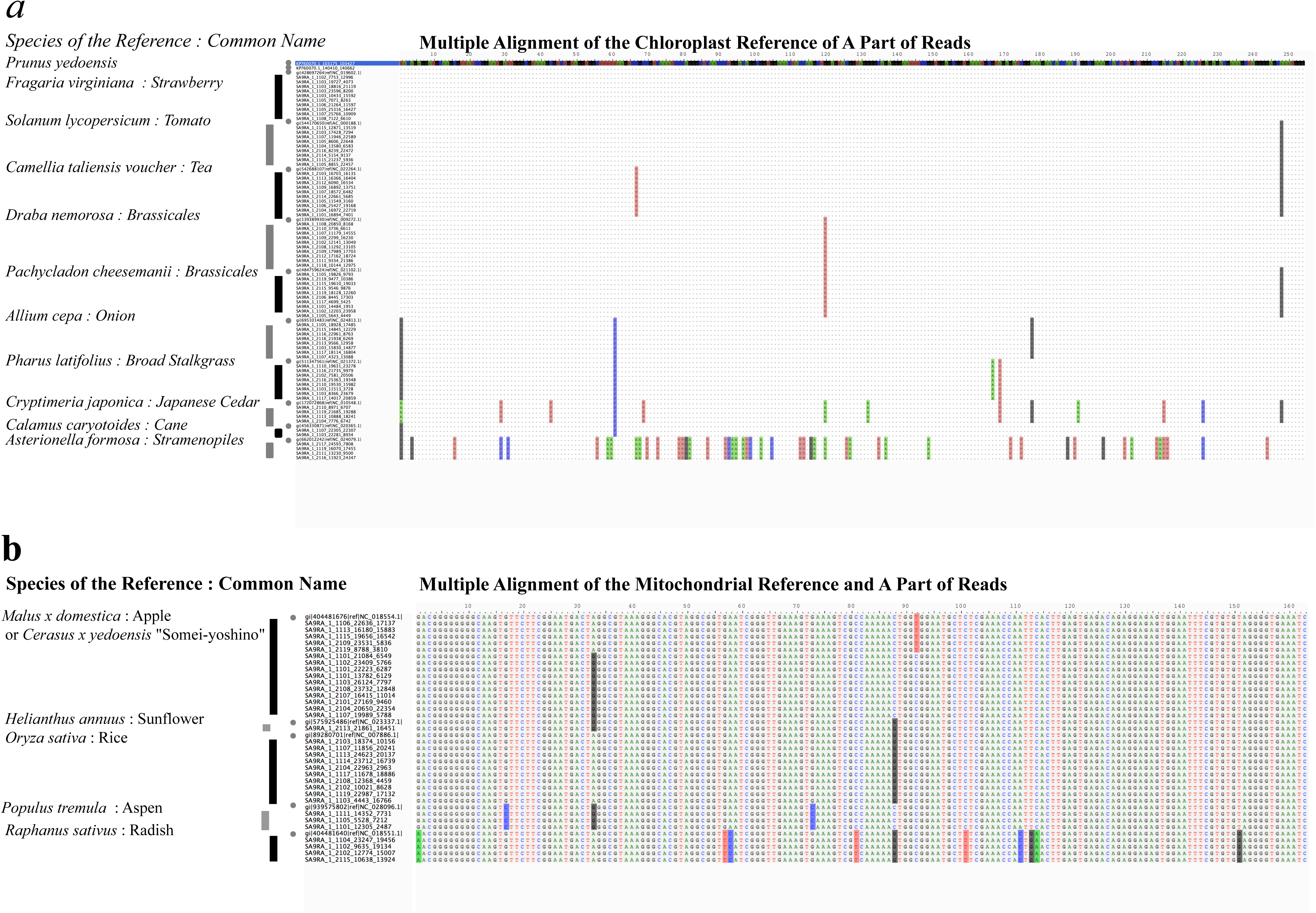
Multiple sequence alignments of arbitrarily selected. **a** 75 reads and 12 reference chloroplast sequences and **b** 33 reads and 5 reference mitochondrial sequences. The gray circles on the left side of the alignment indicate the reference sequences. The black and gray bars on the left side of the alignment indicate the reads sequenced in this research. The reference chloroplast sequences are those of flowering cherry (*Prunus yedoensis*: KP760070.1, 103175 bp to 103427 bp), strawberry (*Fragaria virginiata*: NC_019602.1, 138515 bp to 138767 bp), tomato (*Solanum lycopersicum*: AC_000188.1, 138634 bp to 138886 bp), tea (*Camellia taliensis* voucher: NC_022264.1, 139849 bp to 140101 bp), Brassicales (*Draba nemorosa*: NC_009272.1 and *Pachycladon cheesemanii*: NC_021102.1, 137177 bp to 136925 bp), onion (*Allium cepa*: NC_024813.1, 135931 bp to 136183 bp), broad stalkgrass (*Pharus latifolius*: NC_021372.1, 128467 bp to 128718 bp), Japanese cedar (*Cryptimeria japonica*: NC_010548.1, 107080 bp to 106828 bp), cane (*Calamus caryotoides*: NC_020365, 139357 bp to 139608 bp), and Stramenopile (*Asterionella formosa*: NC_024079.1, 117552 bp to 117800 bp). The mitochondrial sequences are those of apple (*Malus × domestica*: NC_018554.1, 275850 bp to 276100 bp), sunflower (*Helianthus annuus*: NC_023337.1, 141330 bp to 141580 bp), rice (*Oryza sativa*: NC_007886.1, 415532 bp to 415782 bp), aspen (*Populus tremula*; NC_028096.1, 319579 bp to 319829 bp), radish (*Raphanus sativus*: NC_018551.1, 118627 bp to 118877 bp). Every reference sequence was hit by multiple reads at the 100% identity. The sequence alignment indicates that the species detected by BLASTN are actually present on the petal surfaces of the samples and are not a consequence of sequencing errors.

Two other sequences were identified in the BLASTN search regardless of sampling sites, although their read numbers varied among samples: chloroplast DNAs of *Cryptomeria japonica* (Japanese cedar) and *Calamus caryotoides* were observed in 142 and 337 samples, respectively (Fig. 3). *Cryptomeria japonica* is widely distributed across Japan except in Hokkaido and Okinawa (the farthest north and south prefectures), and its wind-pollinated flowers are a serious source of pollen allergies during February–April in Japan. It is likely that *C. japonica* pollen was present on the petals of *C. × yedoensis* flowers and detected as environmental DNA. Likewise, we assumed that the reads identified as *Calamus caryotoides* chloroplast DNA may have originated from a related species that is widely distributed across Japan. The detection of DNA of other plant species suggests that the post-blossoming petal surface is a promising target to reveal dynamics of environmental DNA in nature. A multiple sequence alignment revealed sequence variation among samples collected at different sites (Fig. 4), which may reflect biogeographical structures within each species. With the exception of several Actinobacterial and Proteobacterial reads, we could not obtain a finer resolution on sequences that did not match to organelle DNA, although a rough analysis indicated presence of diverse bacterial species on the petals (Fig. 5).

**Fig. 5.**
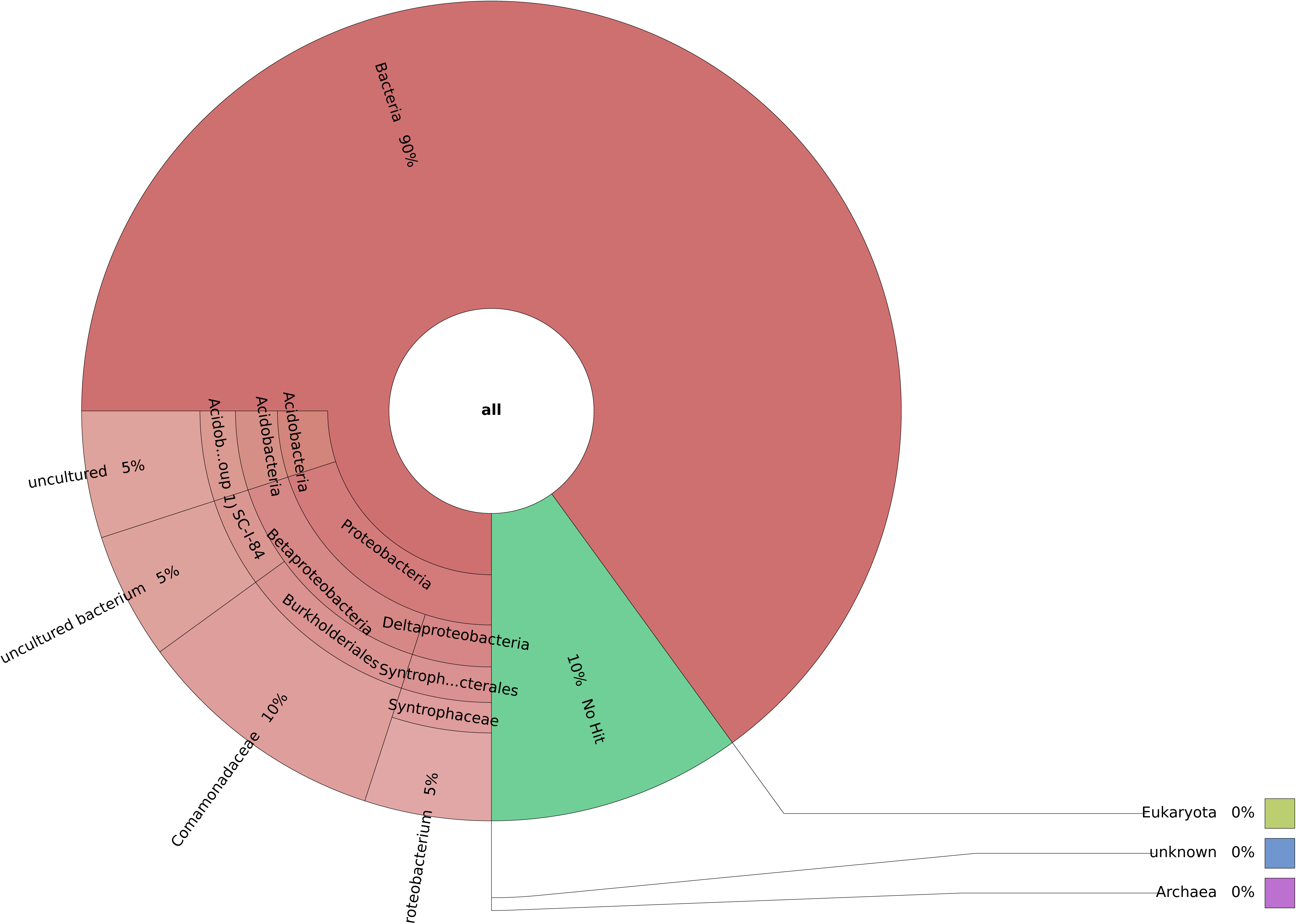
The taxonomic ratio for reads from sample ID 000612 based on BLASTN top hits against the SILVA database. Reads assigned to chloroplast and mitochondria were removed. Visualization was performed using Krona (Ondov et al. 2011). Since the numbers of reads assigned to bacteria were deficient in most samples, improvement of the method of sampling and sequencing may be required to analyze bacterial DNA on petal surfaces.

This is the first large-scale sampling of environmental DNA on petal surfaces from a synchronously flowering species. One of the aims of this study was to use sequencing data to identify the source of environmental DNA on petal surfaces, regardless of their short-term exposure to the environment. We confirmed the presence of mitochondrial and chloroplast DNAs from multiple plant species on the petal surfaces of flowering cherry. On the other hand, identification of the source of bacterial DNAs may require different approaches to sampling and sequencing. With regard to our second aim, we clearly demonstrated the potential value of crowdsourcing for data collection by successfully obtaining numerous samples in a short time frame. As sequencing costs continue to decrease rapidly, we envision the combination of crowdsourced sampling and high-throughput sequencing as a powerful approach to uncovering the dynamics of environmental DNA (Afshinnekoo et al. 2015).

## Acknowledgements

We thank all the Ohanami Project collaborators (Supplementary File 7) for their help with sampling. We also thank the NGS Field 4th Meeting organizers who encouraged us to publish this manuscript. We thank Dr. Hiroshi Mori of National Institute of Genetics for helpful comments. We would also like to show our gratitude to Halna Tsunekawa for graphical design of the project website and the logo.

Computations were partially performed on the NIG supercomputer at ROIS National Institute of Genetics.

## Author contributions

Tazro Ohta, Takeshi Kawashima, Yukuto Sato, Riu Yamashita, Kazuharu Arakawa, and Wataru Iwasaki led the project. Natsuko Shinozaki, Yukuto Sato, and Riu Yamashita performed library construction and sequencing. Tazro Ohta, Takeshi Kawashima, Akito Dobashi, Satoshi Hiraoka, Tatsuhiko Hoshino, Keiichi Kanno, Takafumi Kataoka, Shuichi Kawashima, Motomu Matsui, Wataru Nemoto, Suguru Nishijima, Natsuki Suganuma, Haruo Suzuki, Y-h. Taguchi, Yoichi Takenaka, Yosuke Tanigawa, Momoka Tsuneyoshi, Kazutoshi Yoshitake, Yukuto Sato, and Kazuharu Arakawa performed data analysis. Tazro Ohta developed the data management system and submitted data to the public repository. Takeshi Kawashima, Tazro Ohta, and Wataru Iwasaki wrote the manuscript.

## Legends for Electronic Supplementary Material

**Supplementary File 1.** Sampling protocol handout for participants. The instructions in the sampling protocol were provided to participants in the text and embedded photographs. The instructions also state the protocol for storage and transfer of the collected samples.

**Supplementary File 2.** The histograms of sample metadata. (A) Sampling dates, (B) days since blooming, (C) air temperature, (D) rainfall amount on the sampling date.

The sampling dates and days since blooming were provided by sample submitters, while the meteorological data were retrieved from the website of Japan Meteorological Agency.

**Supplementary File 3.** The complete table of sample metadata. The meteorological data were retrieved from the website of Japan Meteorological Agency.

**Supplementary File 4.** Details of the primers used for the DNA sequencing. Following the 1st PCR, the 2nd PCR was performed to add dual-index tag sequences (D5 and D7 series; Illumina) and then to add flow cell binding sites.

**Supplementary File 5.** Index sequences used for the 1st sequence run. The data submitted to the DDBJ Sequence Read Archive are already demultiplexed and do not contain the index sequences.

**Supplementary File 6.** Index sequences used for the 2nd sequence run. The data submitted to the DDBJ Sequence Read Archive are already demultiplexed and do not contain the index sequences.

**Supplementary File 7.** The member list of Ohanami Project collaborators. All those people have kindly helped the project to succeed this unique crowdsourcing sample collection across the country.

